# Expression and purification of a functional heteromeric GABA_A_ receptor for structural studies

**DOI:** 10.1101/318832

**Authors:** Derek P Claxton, Eric Gouaux

## Abstract

The multi-subunit GABA-gated chloride channels of the Cys-loop receptor family, known as GABA_A_ receptors, function as the primary gatekeepers of fast inhibitory neurotransmission in the central nervous system. In addition to their role in controlling synaptic tone, these receptors are the targets of a vast array of therapeutic compounds that potentiate channel gating. Importantly, functional activity and pharmacological efficacy of GABA_A_ receptors is coupled directly to the subunit composition. However, the absence of high resolution structural information precludes an explicit determination of the molecular mechanism of ligand binding to ion channel gating and modulation. Efforts to obtain this data are hindered largely by the lack of heterologous expression and purification protocols for high expressing receptor constructs. To address this issue, we describe a unique approach to identify bona fide functional GABA_A_ receptor subunit combinations by using the *Xenopus* oocyte as an expression host in combination with fluorescence detection size exclusion chromatography. The results demonstrate that formation of a defined pentameric species is dependent on subunit composition. Furthermore, receptor subunits can tolerate large truncations in non-conserved M3/M4 cytoplasmic loop, although removal of N-linked glycosylation sites is negatively correlated with expression level. Additionally, we report methods to improve GABA_A_ receptor expression in mammalian cell culture that employ recombinant baculovirus transduction. From these methods we have identified a well-behaving minimal functional construct for the α1/β1 GABA_A_ receptor subtype that can be purified in milligram quantities while retaining high affinity agonist binding activity.

## Introduction

The magnitude and duration of excitatory signaling events in the central nervous system is controlled by inhibitory neurotransmission mediated primarily by the opening of pentameric ligand-gated ion channels of the Cys-loop receptor family. GABA (γ-aminobutyric acid) functions as the primary inhibitory chemical messenger and elicits membrane hyperpolarization through rapid diffusion of chloride on the millisecond timescale upon binding to cognate GABA_A_ receptors [1,2]. These ion channels are also the targets of endogenous neurosteroids [3,4] and pharmacological compounds that possess sedative and calming properties through receptor modulation, which defines the clinical basis of general anesthetics and anxiolytics [5-7]. The critical role of these receptors in suppressing excitatory inputs is emphasized further by the strong correlation between reduced GABA_A_ receptor activity and seizure disorders that vary in severity, including the most debilitating epileptic encephalopathies [8-10].

The potential to alter channel gating by allosteric modulators suggests that GABA_A_ receptors are excellent candidates for targeted neurological therapies. As an example, the benzodiazepine drug class has been employed as the choice therapy for controlling epilepsy [11], anxiety [12] and sleep disorders [13] by directly potentiating GABA_A_ receptor activity. However, parallel to the broad therapeutic benefits of benzodiazepines is the presentation of adverse side effects, such as amnesia and addiction [7], as a result of long term use. Importantly, the explicit molecular mechanism of drug modulation on GABA_A_ receptors is not known, although the serendipitous discovery of benzodiazepines was made more than 50 years ago [14].

The chief contributor to an incomplete understanding of GABA_A_ receptor function and modulation is the lack of atomic scale structural models that define ligand binding sites and conformational states. Presently, structural and functional studies are interpreted from homology models generated from bacterial homologs [15-18] or related Cys-loop receptors, such as the nicotinic acetylcholine receptor [19-21] or the glutamate-gated chloride channel [22-24]. In general, all Cys-loop receptors share a common core architecture defined by formation of five homologous subunits arranged about a central ion-conducting pore as observed in structures of homo-pentameric channels [23,25,26]. Each subunit consists of a large extracellular amino-terminal domain connected to a four-helix bundle transmembrane domain that contributes residues for ion selectivity. The orthosteric binding site is formed at the interface of adjacent subunits in the amino-terminal domain. Coupling between agonist binding and channel gating is facilitated by an interaction network composed of residues in transmembrane and extracellular domain loops, including the characteristic disulfide [27].

Despite the similarities between all Cys-loop receptors, homology models may not capture the subtle features of GABA_A_ receptors, in part because there are a large number of heteromeric receptor assemblies with discrete ion channel and pharmacological properties [28]. So far 19 subunits have been identified and separated into classes and isoforms depending on the primary sequence: α(1-6), β(1-3), γ(1-3), δ, ε, θ, π and ρ(1-3). Although formation of homo-pentamers has been reported from *in vitro* studies [25], recapitulation of *in vivo* gating and pharmacology requires two or more subunits. Even though random assembly is unlikely [29], the molecular determinants that drive assembly of specific combinations in defined subunit stoichiometries and organization are not entirely known. Assembly signals encoded within the subunit primary sequence may play a role in limiting receptor subtype diversity [30]. Notably, subunit expression patterns suggest that unique properties of specific subunit combinations are fine tuned to regulate either phasic or tonic inhibition [28]. The most widely distributed receptor combination in adult neurons is the α1/β2/γ2 subtype in a predicted 2α:2β:1γ subunit ratio [31-34]. However, receptors composed of only α/β subunits (2α:3β), which form channels that are strongly antagonized by Zn^2+^, may play a role during embryonic development [35-39]. Importantly, incorporation of the γ subunit confers sensitivity to benzodiazepines and the derived clinical benefit is tied to the adjacent α subunit isoform [7,40]. Furthermore, the promiscuity of classical, non-selective benzodiazepines, such as diazepam, contributes not only to the therapeutic benefits, but also to the undesired side effects.

These observations emphasize the need for accurate, experimentally-derived molecular models of GABA_A_ receptors that define subunit-specific interactions and that link ligand binding to gating activity to identify mechanistic differences between subtypes and facilitate structure-based drug design. As a prerequisite, identification of efficient heterologous expression and isolation methodologies are required to generate sufficient quantities of receptor for structural interrogation via X-ray diffraction or single particle cryo-electron microscopy. Development of such methods is not trivial for eukaryotic membrane proteins because functional expression may be dependent on the host system, require co-factors or posttranslational modifications [41,42]. Here, we outline a novel approach for identifying and characterizing a well-behaving heteromeric GABA_A_ receptor composed of α/β subunits. This approach combines the robust translation efficiency of *Xenopus* oocytes for screening subunit constructs with technological advances for large-scale expression in mammalian cell culture using a modified baculovirus delivery system [43]. We show that the α1/β1 receptor can be expressed and purified in milligram quantities while retaining ligand binding activity. These results set the stage for detailed structural investigation of heteromeric GABA_A_ receptors and provide a foundation for pursuing other multi-subunit Cys-loop receptor assemblies.

## Materials and Methods

### Construction of vectors, generation of fluorescent protein fusions and mutagenesis

The full-length rat GABA_A_ receptor subunit isoforms α1 (GeneID 29705), β2 (GeneID 25451) and γ2S (GeneID 29709) with the native signal sequence were obtained in the pGEM vector as a gift from Dr. David S. Weiss. GluClα (Gene ID 180086) was also inserted into the pGEM vector. This vector contains a T7 RNA polymerase promoter site for *in vitro* RNA transcription. For expression in insect cells, individual subunits with a leading Kozak sequence were cloned into the pFastBac1 vector where expression is driven by the polyhedron (PH) promoter, or into the pFastBac Dual vector in which case a single virus particle contains both α1 (PH promoter) and β2 (p10 promoter) subunit genes. For mammalian cells, all genes were cloned individually into a novel vector, pEGBacMam [43], where expression is controlled by the human cytomegalovirus (CMV) promoter.

Identification of the signal peptide for each subunit was made using the SignalP 4.1 server [44]. Residue numbering is based on the mature subunit sequence following cleavage of the predicted signal peptide. The fluorescent protein EGFP was inserted into GluClα in the M3/M4 loop as described previously [45]. EGFP was ligated to the M3/M4 loop of the α1 subunit between V372 and K373 using an in-frame non-native Asc1 restriction site (GGGCGCGCC), which adds a three residue linker (G-R-A) on each side of the fluorescent protein to the native receptor sequence. Likewise, mKalama was inserted into the β2 subunit M3/M4 loop between H397 and V398 also using an Asc1 restriction site. To facilitate M3/M4 loop truncations (“LT”) in α1, an amino-terminal fluorescent protein fusion of the α1 subunit was made by inserting GFPuv after the signal peptide with a 24 residue polypeptide linker (S-S-S-N-N-N-N-N-N-N-N-N-N-L-G-T-S-G-L-V-P-R-G-S) containing a thrombin protease site. The α1-LT construct was generated by replacing M3/M4 cytoplasmic loop residues Y314-P381 with Thr for a final predicted M3/M4 linker of R-G-T. Likewise, β2-LT was generated by replacing R309-I414 with Gly for a final predicted M3/M4 linker of G-G-T. The β1-LT subunit gene (GeneID 25450) was synthesized and cloned into pEGBacMam by Bio Basic Inc with residues P310-K413 replaced with Gly-Thr for a final predicted M3/M4 linker of G-G-T. Incorporation of fluorescent protein fusions and receptor mutations were made using standard PCR procedures and confirmed by DNA sequencing.

### RNA synthesis, oocyte injection, microscopy, FSEC and functional analysis

Circular plasmid DNA in the pGEM vector was linearized with NheI restriction endonuclease followed by *in vitro* synthesis of capped mRNA using the mMESSAGE mMACHINE T7 Ultra kit from Ambion. After polyadenylation, mRNA was precipitated with LiCl, washed with 70% ethanol and re-suspended with DEPC-treated water. Concentration of mRNA was determined by absorbance at 260 nm and the quality was judged by gel electrophoresis. Defolliculated *Xenopus* oocytes (stage V-VI) were injected with 25-50 ng of total mRNA mixed in defined ratios to obtain expression of specific receptor subtypes. Oocytes were incubated at 16-18 °C in ND-96 solution (96 mM NaCl, 2 mM KCl, 1 mM MgCl_2_, 1.8 mM CaCl_2_, 5 mM Hepes pH 7.5) supplemented with 250 μg/mL Amikacin for three to five days. Eight to 12 oocytes were mechanically disrupted and solubilized in 150 μL TBS pH 7.4 including 40 mM n-dodecyl-β-D-maltopyranoside detergent (C_12_M) and 2 mM PMSF for 45 minutes with gentle mixing. The sample was then centrifuged at 186,000 rcf for 40 minutes and the supernatant loaded onto a Superose 6 10/300 GL column (GE Healthcare) for analysis by fluorescence detection size exclusion chromatography (FSEC) monitoring EGFP (510 nm) [46]. Receptor surface expression was observed using a Zeiss LSM710 laser scanning confocal microscope. Resonance energy transfer studies employed the high intensity laser (488 nm) to photobleach EGFP and subsequently monitor mKalama fluorescence from 410-486 nm. For two electrode voltage clamp electrophysiology (TEVC) experiments, 5 ng of total mRNA was injected into oocytes. After one to three days incubation, oocytes were voltage clamped to either −30 mV or −60mV and perfused with 0.1 or 1mM GABA in the absence or presence of 10μM zinc in ND-96 buffer.

### Transient Transfection

Approximately 1×10^6^ TSA201 cells (ATCC) were allowed to attach to a 35mm dish bathing in DMEM media supplemented with 10% FBS. Subunit plasmid DNA (2-3 μg total) in pEGBacMam were incubated with Lipfectamine 2000 reagent in OPTIMEM media for 20 minutes and then added dropwise to the cells showing 80-90% confluence. After 24 hours at 37 °C, the transfection mixture was replaced with fresh DMEM supplemented with 10% FBS and 5mM sodium butyrate. The cells were incubated an additional 30 hours at 37 °C prior to harvesting for FSEC analysis.

### Recombinant baculovirus production

Baculovirus was generated as described previously [47,48]. Briefly, subunit constructs in pFastBac1, pFastBac Dual or pEGBacMam vectors were transformed into DH10Bac competent cells for site-specific transposition of the expression cassette into bacmid. Purified bacmid was transfected into 1×10^6^ cells Sf9 cells (Invitrogen^TM^) in a 35 mm dish cultured in serum free Sf-900 III SFM medium using Cellfectin II reagent and P1 virus was harvested four days later. Virus was further amplified in Sf9 cells and P2 virus harvested after four days. Viruses were stored at 4 °C in the dark supplemented with 2% FBS. Virus titer was determined using the Sf9 Easy Titer cell line and the end-point dilution assay [49] or flow cytometry by using the ViroCyt Virus Counter 2100 [50].

### Insect cell expression

P2 virus was used to infect Sf9 or High Five cells (Invitrogen^TM^) at a density of 1.5-3×10^6^ cells/mL cultured in serum free Sf-900 III SFM or Express Five medium using different absolute and relative (subunit ratio) MOI. Infected cells were incubated with shaking at either 27 °C for the duration of the expression trial, or shifted to 20 °C after an initial 18 hours at 27 °C. Each day for five days, 1 mL of cells were pelleted by centrifugation and frozen at −80 °C. The cells were solubilized in TBS pH 7.4 with detergent supplemented with a protease inhibitor cocktail for FSEC analysis as described for oocytes.

### Expression and purification from mammalian cells

High titer P2 virus was used to infect TSA201 or HEK293 GnTI^−^ suspension cells (ATCC) at a density of 1.5-3×10^6^ cells/mL cultured in Gibco Freestyle 293 Expression medium supplemented with 2% FBS and placed in a humidity- and CO_2_-controlled incubator. The total volume of virus added was less than 10% of the culture volume in all cases. After 12 hours at 37 °C, sodium butyrate was added to 10 mM and the culture shifted to 30 °C. After 60-72 hours post-infection, cultures were harvested by centrifugation and processed by sonication in the presence of a protease inhibitor cocktail. Membranes were isolated by ultracentrifugation, mechanically homogenized and solubilized with TBS pH 8.0 including 40 mM C_12_M. The solubilized receptor was harvested by ultracentrifugation and mixed with ClonTech Talon Co^2+^ affinity resin for three to four hours. The resin was washed with a low concentration of imidazole and then eluted with 250 mM imidazol buffer including 1 mM C_12_M or 0.3 mM lauryl maltose neopentyl glycol (L-MNG) detergent. Fractions containing the receptor were pooled and concentrated using a 100 kDa MWCO filter to 0.5mL for purification over a Superose6 10/300 GL column. Receptor purity was assessed by SDS-PAGE and coomassie blue staining using a 12.5% Tris-HCl gel. LC-MS was employed to confirm the identity of subunit bands on the gel. Receptor concentration was determined by absorbance at 280 nm assuming an average 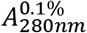 determined by the following equation:

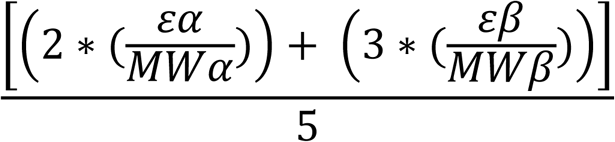
Here we assume a subunit ratio of 2:3, α:β, and calculated molecular weights (MW) and molar absorption coefficients (*ε*) for each mature subunit.

### Radioligand receptor binding

Titration of purified receptor using 500 nM muscimol (10% hot, Perkin Elmer) and 1 mg/mL YSi Copper His tag SPA beads suggested a capacity of 100 nM receptor (~100 pmol) per mg of beads. For saturation binding and competition assays, 5 nM purified receptor was mixed with ^3^H-muscimol (1:10 dilution with cold muscimol, yielding a specific activity of 2-4 Ci/mmol) and 1mg/mL SPA beads in 20 mM Hepes pH 7.35, 150 mM NaCl, 0.3 mM L-MNG in 100 uL final reaction volume. Muscimol was titrated from 3-400 nM to determine *K*_D_, or held constant at 100 nM to determine *K*_i (GABA)_. Counts were recorded on a MicroBeta TriLux luminescence counter from Perkin Elmer. Specific binding was measured with 1 mM GABA as a cold competitor. Each data point was repeated in triplicate for two separate trials and the standard deviations determined. Muscimol *K*_D_ was determined from nonlinear least squares fit of specific binding assuming a single site binding model. The IC_50_ of GABA was determined from fitting the data points of the competition experiment with the Hill equation, which was then used to determine *K*_i_ using the following equation:

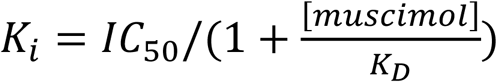

All fits were performed with the program Origin (OriginLab Corp).

## Results

### Screening methodology of Cys-loop receptor subunit constructs

Historically, *Xenopus* oocytes have proven a convenient vehicle for heterologous gene expression to assess exogenous ion channel properties by electrophysiological methods because they possess the machinery for gene expression but have low endogenous receptor activities [51,52]. We have taken advantage of the robust gene expression in oocytes as a tractable platform for screening DNA constructs of the multi-subunit Cys-loop receptors. For our studies, genes were designed as fusions with GFP variants and delivered into the cytoplasm by microinjecting mature oocytes with up to 50 ng of total capped RNA transcripts synthesized *in vitro*. After three to five days at 16 °C, a small batch of oocytes (8-12) expressing the same construct was solubilized in detergent for analysis by fluorescence detection size exclusion chromatography or FSEC [46].

As a benchmark, oocytes were injected with either 20 ng or 50 ng of RNA encoding the well-behaved α subunit of the glutamate-gated chloride channel (GluCl) from *C. elegans*, a homomeric Cys-loop receptor [23]. The GluCl construct was designed as a fusion with EGFP within the large cytoplasmic M3/M4 loop, which has been shown to not disrupt channel function [45] (Fig 1A). After four days, EGFP fluorescence could be observed at the oocyte surface relative to an uninjected control using a laser scanning confocal microscope, consistent with expression of the receptor (Fig 1B). Oocyte solubilization with the mild detergent n-dodecyl-β-D-maltopyranoside (C_12_M) and EGFP-FSEC analysis allowed detection of the receptor peak eluting at the approximate time for the intact pentamer based on molecular weight standards and demonstrated sensitivity to the dosage of RNA injected (Fig 1C). Solubilization of uninjected oocytes outlined peaks that are likely to arise from the oocyte itself.

**Fig 1.**
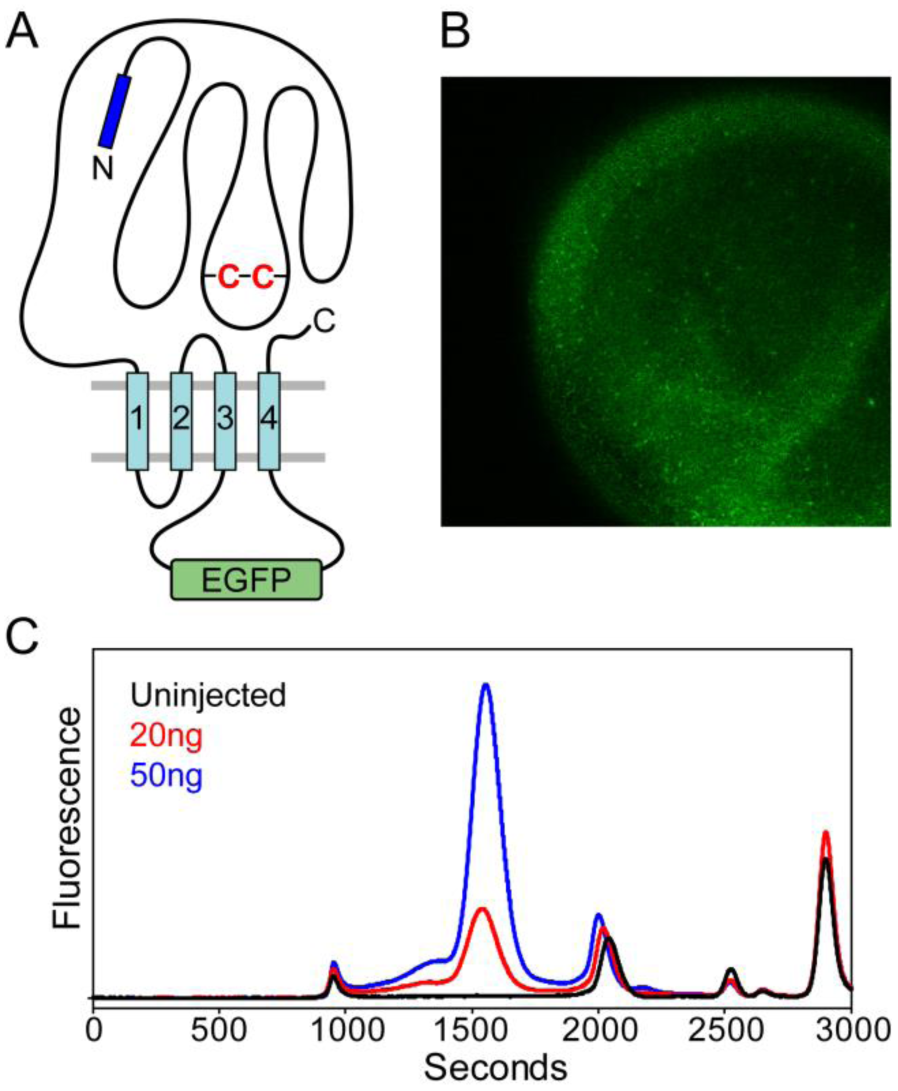
Expression of GluClα-EGFP in oocytes. (A) Cartoon design of EGFP fusion to the M3/M4 loop of GluCl. (B) Four days after injecting 50 ng of synthetic mRNA, receptor expression is visualized by EGFP fluorescence at the oocyte surface by confocal microscopy. (C) Detergent (C_12_M) solubilization of oocytes and FSEC analysis captures a monodisperse elution profile at ~1550 seconds that is consistent with formation of pentameric receptor. Receptor expression levels are sensitive to the total concentration of mRNA injected.

### Oocyte expression of GABA_A_ receptor subunits

The propensity for GABA_A_ receptor subunits to form homo- or heteromeric channels was investigated by injecting oocytes with the full-length rat α1 isoform as a unitary subunit or in combination with full-length β2 or γ2S subunits or with all three subunits. Analogous to GluCl in Fig 1A, the α1 subunit was expressed initially with an EGFP fluorescent protein fusion within the non-conserved M3/M4 loop (α1-EGFP), whereas the other subunits were expressed as wild type sequences. For expression of heteromeric assemblies, synthetic RNA transcripts were combined in defined ratios while keeping the total RNA injected constant (25-50 ng).

Three days post-injection, oocytes were solubilized in C_12_M and analyzed by EGFP fluorescence for expression and profile homogeneity by FSEC. Although the α1 subunit alone demonstrated oligomerization near the elution position of a pentamer, peak shape is consistent with a heterogeneous entity (Fig 2A). Other peaks in the FSEC trace suggest the presence of lower order oligomeric species (arrow, Fig 2A). Interestingly, co-injection with the β2 subunit increases overall expression and monodispersity of the primary peak and a reduction in the population of other oligomeric states, suggesting a change in subunit composition coincides with pentamer stabilization (Fig 2A). Notably, similar profiles are observed when α1-EGFP or α1-EGFP/β2 is transiently transfected into adherent TSA201 cells, although absolute yields are lower relative to oocytes (Fig 2B).

**Fig 2.**
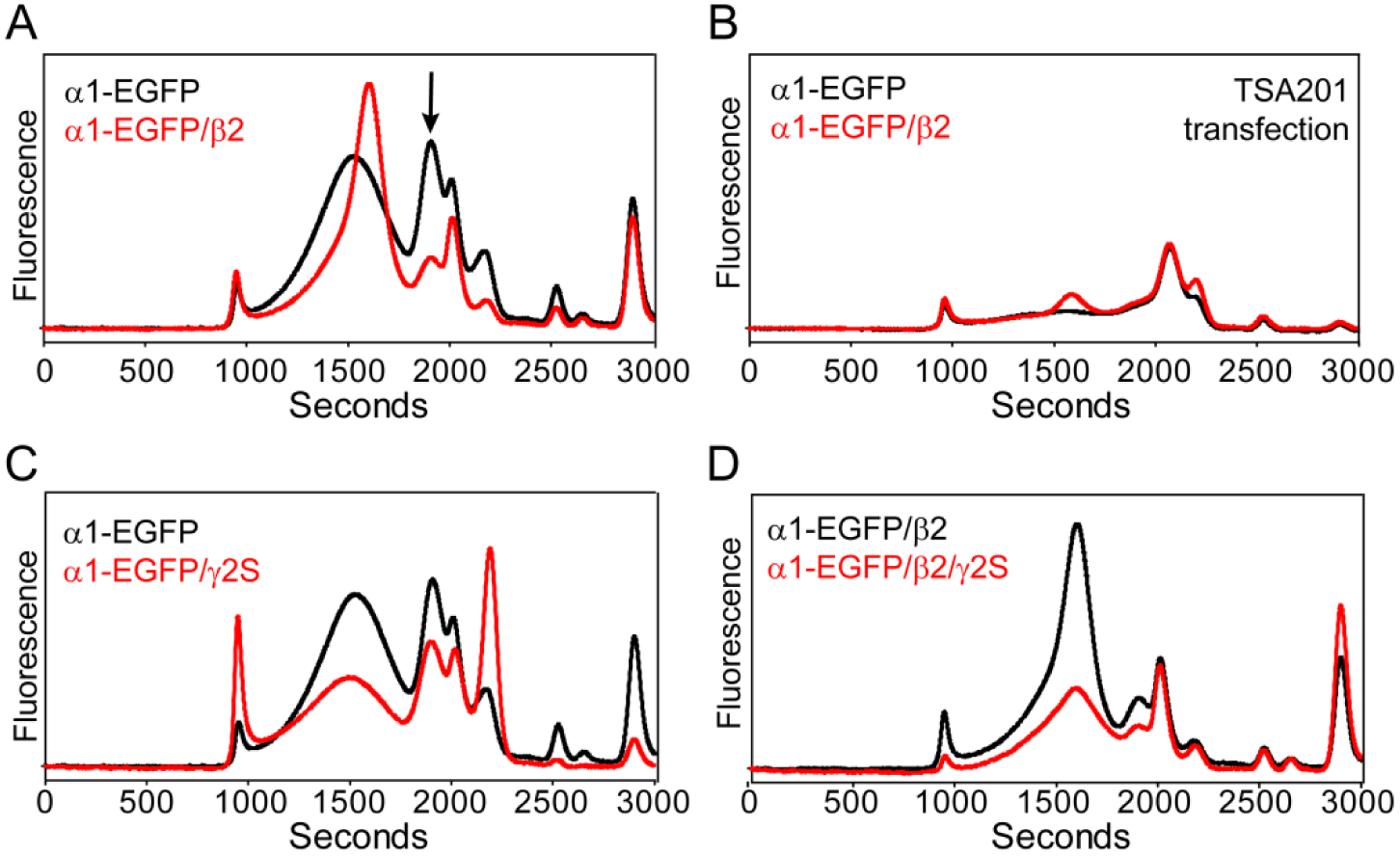
Expression of GABA_A_ receptors in oocytes. (A) The α1/β2 receptor demonstrates increased expression levels and homogeneity relative to the α1 subunit alone. The arrow indicates a population reduction of smaller oligomers in the presence of the β2 subunit. (B) Similar profiles are obtained from transient transfection of TSA201 cells incubated at 37 °C, although expression levels are substantially lower than in oocytes. (C) Co-injection of γ2S mRNA with the α1 subunit does not improve receptor profiles. (D) Injection of all three subunits results in attenuated expression levels relative to α1/β2. All FSEC traces are plotted on the same scale.

In contrast, co-injection with the γ2S (short splice variant) subunit did not change the profile relative to the α1 subunit alone (Fig 2C), which may indicate that α1/γ2S receptors are unlikely to form channels in oocytes. However, we know that addition of the γ2S subunit in the presence of α1/β2 enables formation of the most common tri-heteromeric receptor in the central nervous system [32,33]. Relative to the α1/β2 subunit combination, injection of all three subunits (α1-EGFP/β2/γ2S, 12ng each) resulted in substantially reduced expression levels, which could be due to RNA competition with the available pool of oocyte ribosomes or represent an intrinsic property of α1/β2/γ2 receptors (Fig 2D).

We further investigated the putative association of α1 and β2 subunits by fluorescence resonance energy transfer (FRET) [53]between mKalama, a blue-shifted fluorescent protein, fused to the β2 subunit M3/M4 loop and α1-EGFP using a laser confocal microscope. If FRET occurs, donor (mKalama) fluorescence will be quenched in the presence of the acceptor (EGFP). As shown in Fig 3, photobleaching EGFP at the oocyte surface resulted in an increase in mKalama fluorescence relative to an unbleached control region, suggesting that the fluorescent protein pair is in close enough proximity to undergo FRET and is consistent with the formation of a heteromeric GABA_A_ receptor composed of α1/β2 subunits.

**Fig 3.**
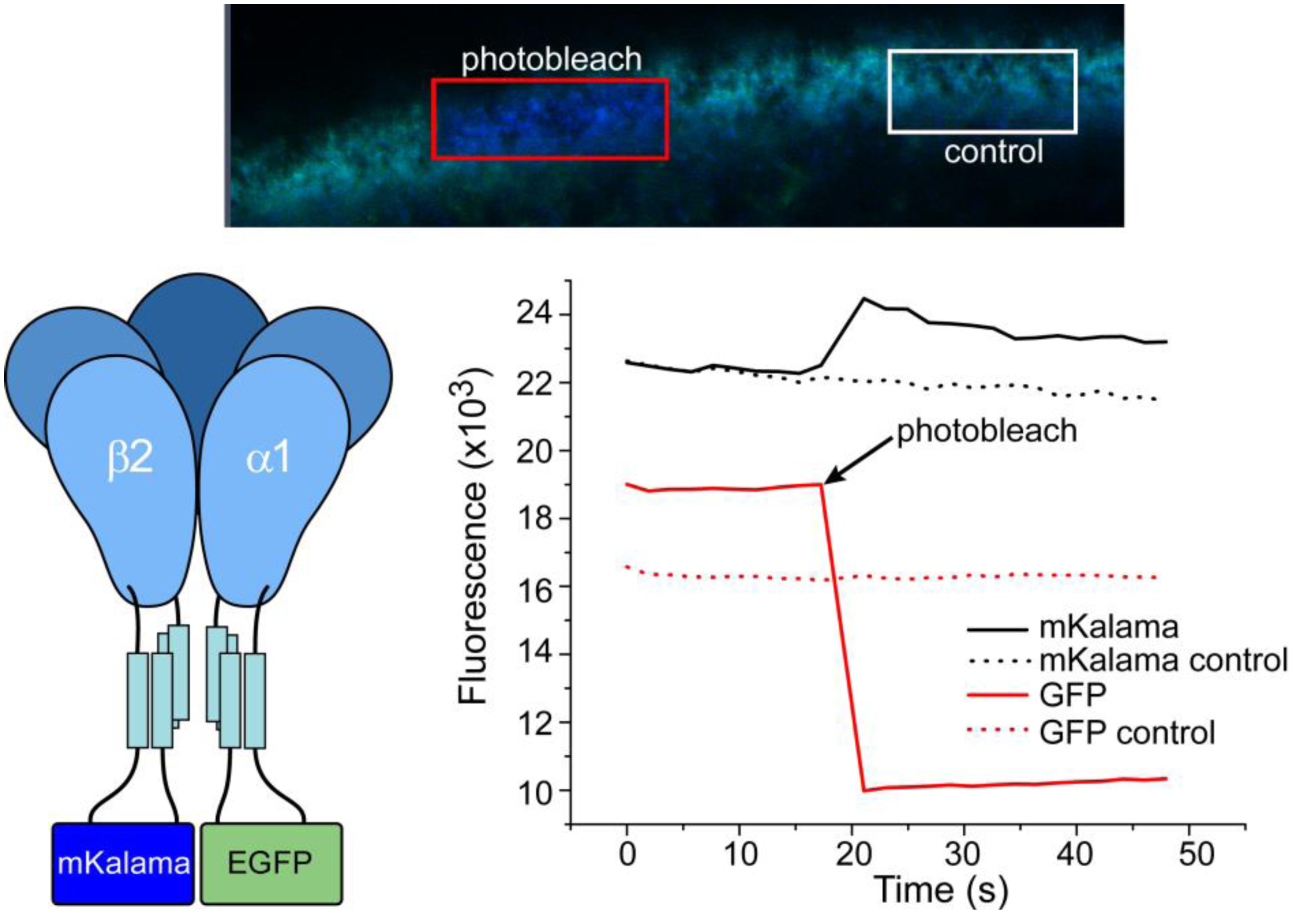
Investigation of α1 and β2 subunit association through FRET. Photobleaching of EGFP fused to the α1 subunit results in increased β2-mKalama fluorescence at the oocyte surface relative to a control region, indicating that the subunits are assembled together to allow the fluorescent proteins to undergo FRET.

Based on preliminary screening of a restricted pool of subunits, the α/β GABA_A_ receptor demonstrated the best expression levels relative to the other tested combinations. To determine a functional α/β construct suitable for downstream large-scale expression, purification and structure determination using X-ray diffraction methods, we employed systematic mutagenesis to trim potentially unstructured polypeptide regions and remove sites of putative chemical modification in both subunits. The mutagenesis was guided by secondary structure prediction algorithms from the PredictProtein server [54] in conjunction with sequence alignments with the homologous GluCl in which a high resolution structure has been solved [23].

### M3/M4 loop truncations

The greatest sequence divergence in alignments of GABA_A_ receptor subunits and of Cys-loop receptors in general is found in the large cytoplasmic loop connecting transmembrane helices three and four (M3/M4 loop). Electron microscopy of the related nicotinic acetylcholine receptor has revealed some secondary structure for this loop, but the majority of the peptide chain remains structurally undefined [19]. Although this loop contains sequence that facilitates protein-protein interactions [55] and single channel conductance [56], macroscopic ion channel properties of Cys-loop receptors persist when the native loop has been removed [57]. In GluCl, the entire 58 residue M3/M4 loop was replaced with the tri-peptide linker Ala-Gly-Thr without altering pentamer formation or channel function [23].

To facilitate biochemical screening of both α and β subunits yet while retaining the native M3/M4 loop, a new fluorescence protein fusion was generated with the cycle 3 GFP variant, GFPuv [58] attached to the N-terminus of the α1 subunit with a 24 amino acid linker and following the predicted signal peptide, designated GFPuv-α1 (Fig 4A). Previous work with the nicotinic acetylcholine receptor indicated that N- or C-terminal GFP fusions did not support proper folding of the chromophore, necessitating placement of GFP in the M3/M4 loop [59]. However, GFPuv is more stable [60] and has been used to study the secretory pathway of class C GPCRs [61]. RNA injection of this fusion construct in oocytes and subsequent FSEC analysis indicated that the GFPuv-α1/β2 receptor expressed with reduced aggregation propensity relative to α1-EGFP/β2 (Fig 4B). Of note, this construct demonstrated higher levels of free fluorescent protein following solubilization as indicated by the arrow in Fig 4B, which is likely a consequence of the long polypeptide linker and protease site between GFPuv and the α1 N-terminus. Nevertheless, the FSEC profile is consistent with a well-behaved receptor, suggesting that the construct is useful for subsequent mutagenesis.

**Fig 4.**
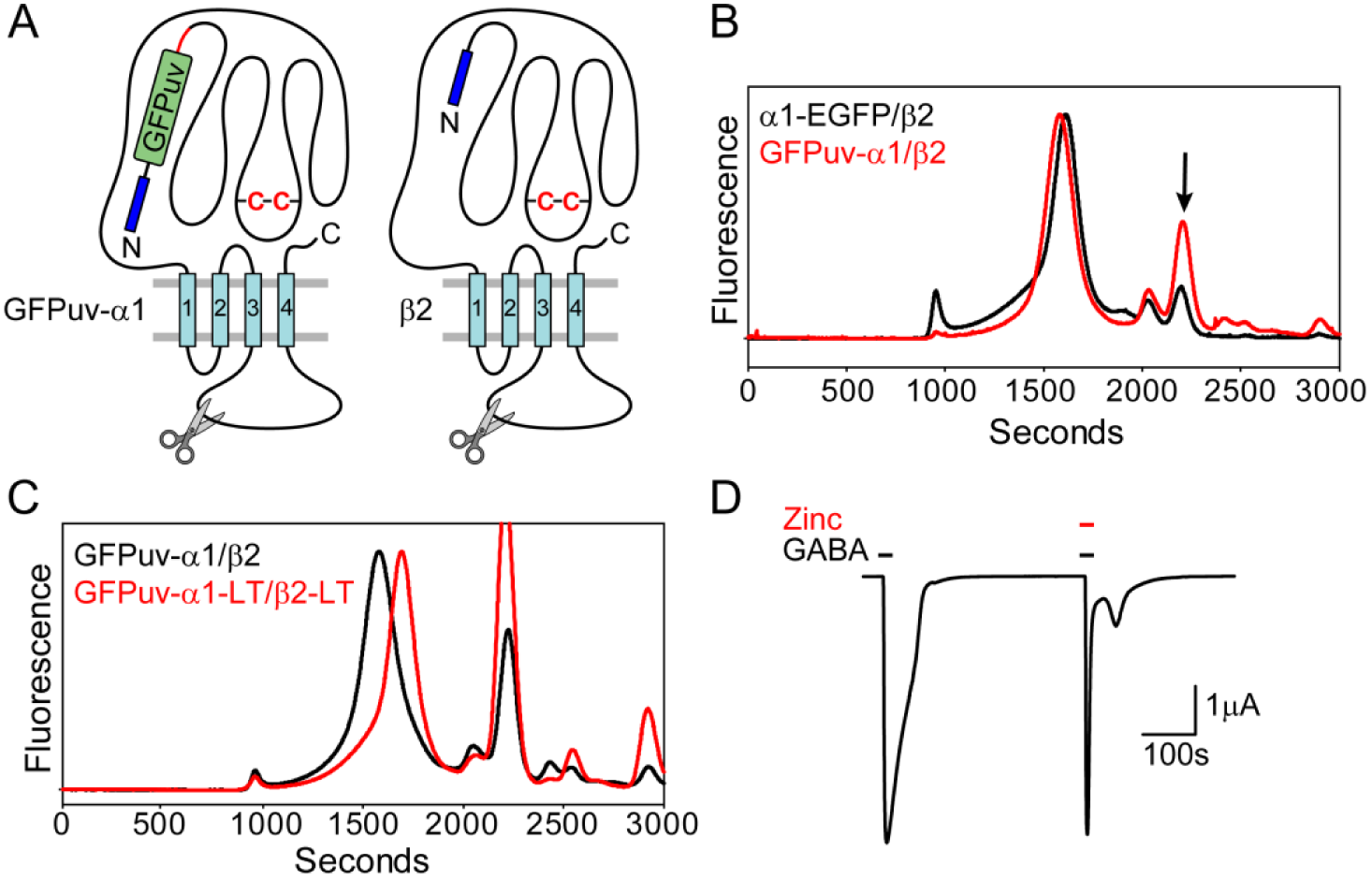
Truncation of the M3/M4 loop does not perturb receptor assembly or function. (A) Cartoon of subunit construct design for α1 and β2 subunits shows locations for GFPuv fusion and loop truncation. A polypeptide linker (red line) connects GFPuv to the N-terminus of the α1 subunit. (B) Expression of GFPuv-α1/β2 increases receptor homogeneity relative to α1-EGFP/β2. However, an increase in cleaved fluorescence signal is observed (arrow). (C) Replacement of the native M3/M4 loop in both subunits with a tri-peptide induces a right shift in the elution profile, consistent with a smaller hydrodynamic radius. (D) Two-electrode voltage clamp of GFPuv-α1-LT/β2-LT demonstrating that the receptor retains gating activity and sensitivity to zinc.

A large truncation of the M3/M4 loop was designed for both subunits, reminiscent of the optimized GluCl construct [23], to investigate changes to the receptor elution profile. In the α1 and β2 subunits, the 70 and 108 residue M3/M4 loops were replaced with Arg-Gly-Thr and Gly-Gly-Thr tripeptide linkers, respectively (defined as “LT”). Combining the loop truncation in both subunits reduced the molecular weight of the receptor by nearly 51 kDa assuming a subunit assembly ratio of 2:3, α1 to β2. This change in mass is reflected in a 100 second right shift in the FSEC elution peak of GFPuv-α1-LT/β2-LT (Fig 4C). We confirmed that this receptor construct retained gating activity by TEVC experiments. A large amplitude current (μA) was elicited from perfusion with 0.1 mM GABA and was also antagonized by Zn^2+^ (Fig 4D).

### N- and C-terminal deletions and removal of glycosylation sites

Construct optimization of GluCl further required deletion of a short carboxy terminal tail and a long stretch of 41 residues in the amino terminus following the signal peptide [23] (Fig 5A-B). In sequence alignments, the α1 carboxy terminus extends well beyond both GluCl and β2 subunits. The last eleven residues in the α1 subunit were removed by mutagenesis. Oocyte expression of this mutant in the GFPuv-α1-LT/β2-LT context did not perturb receptor assembly (Fig 5C). Both GABA_A_ receptor α1 and β2 subunits are predicted to have longer signal peptides relative to GluCl, but shorter amino-termini that precede the predicted α-helix in the extracellular domain. In striking contrast to deletion of the α1 carboxy terminus, deletion of nine residues from the amino-terminus in the α1-EGFP subunit and seven residues in the full length β2 subunit demonstrated substantially reduced expression levels relative to wild type as indicated by lower absolute fluorescence intensity in the FSEC trace (Fig 5D). Combining the deletion constructs nearly abolished expression of the receptor.

**Fig 5.**
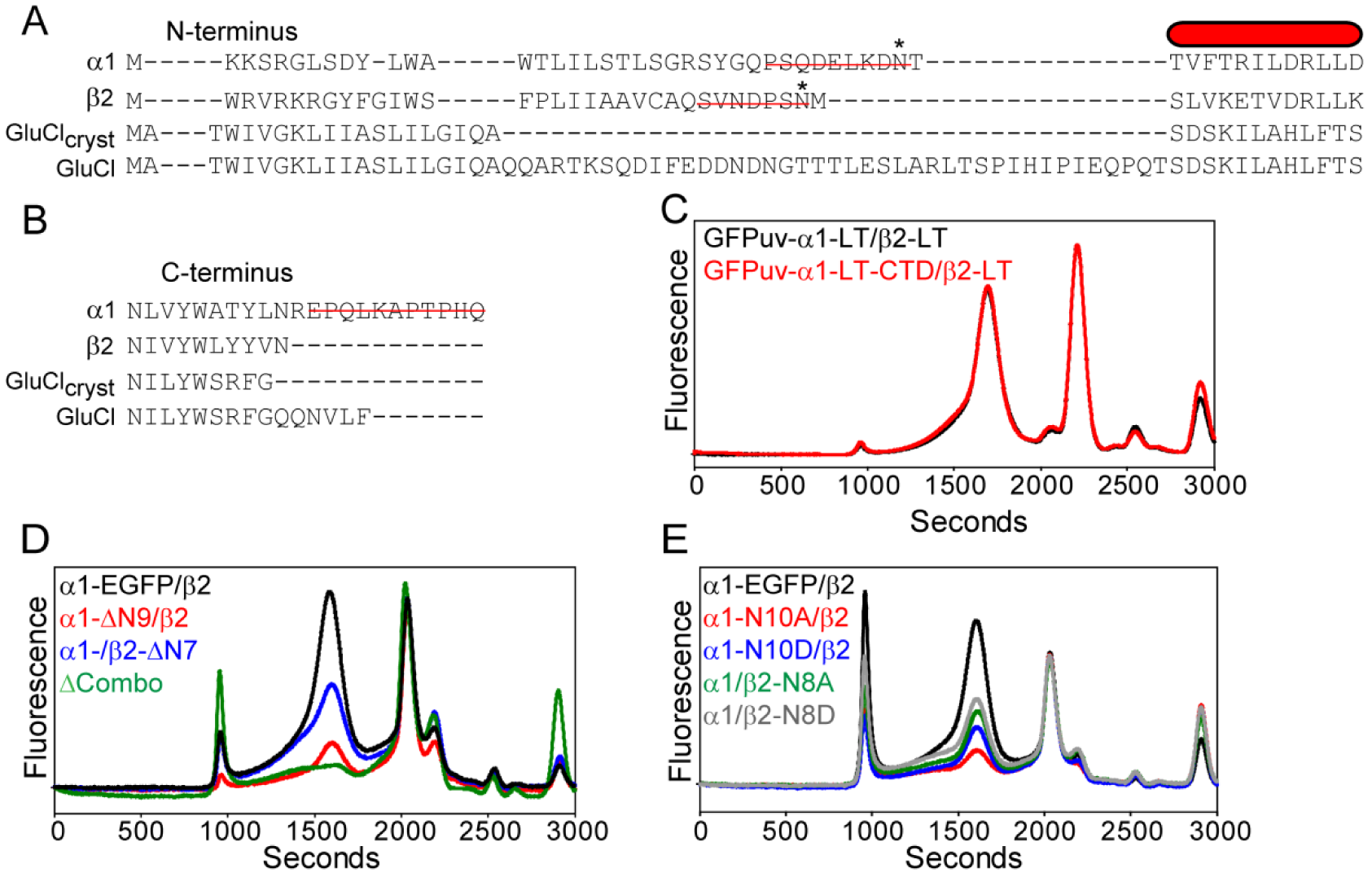
Removal of glycosylation sites on the N-terminus reduces α1/β2 expression. (A) Sequence alignment of the N-terminus for GABA_A_ and GluCl subunits identifying residues for deletion (red line) and predicted sites for glycosylation (*). The red bar represents the first predicted α-helix in the extracellular domain. (B) Sequence alignment of the C-terminus for GABA_A_ and GluCl subunits identifying residues for deletion (red line). (C) Removal of an 11-residue tail from the α1 subunit does not change receptor behavior. (D) Deletion of the N-terminus in either α1 or β2 subunit reduces receptor expression. (E) Site directed mutagenesis of predicted glycosylation sites reduces receptor expression.

Both subunit amino-terminal deletions contain a consensus site for *N*-linked glycosylation (Asn-X-Ser/Thr), which may play a role in surface expression and function of GABA_A_ receptors [62,63]. To test this possibility, each of these sites (α1-N10, β2-N8 in the mature receptor) was mutated to either Asp or Ala in the wild type subunit background and then each subunit was individually examined for impact on receptor expression characteristics. As shown in Fig 5E, even though pentameric receptor was formed, expression levels were attenuated to similar levels as the amino-terminal deletion constructs with mutation of either Asn residue, although mutation in the β2 subunit was more tolerated. This result suggested that prevention of posttranslational modification with N-glycans is the source of diminished expression levels. However, based on these experiments alone, we cannot rule out the potential negative contribution of other deleted residues in the N-terminus to overall expression.

In addition to these sites, the α1 and β2 subunits contain one (N110) and two (N80, N149) other putative glycosylation sites, respectively. The impact of posttranslational modifications at these sites was investigated using the GFPuv-α1-LT/β2-LT construct. Site directed mutagenesis experiments indicated that mutation of both β2 subunit sites lowered expression levels and increased receptor heterogeneity (Fig 6), suggesting that all three glycosylation sites are needed for proper folding and/or targeting. This result is consistent with the observation that mutation of each of the three sites in β2 reduced peak amplitudes and altered gating properties in cultured mammalian cells [63]. However, mutation of N110 in the α1 subunit does not impede receptor expression in oocytes or substantially change monodispersity, which likely indicates that glycosylation at this site is not required for receptor assembly. Although previous studies suggested that mutation of N10, N110 or both to Gln in α1 does not impair channel gating in oocytes [62], bulk expression levels appear to be sensitive to the removal of the N10 glycosylation site based on these results. Overall, the data is consistent with the notion that glycosylation plays a crucial role in receptor maturation.

**Fig 6.**
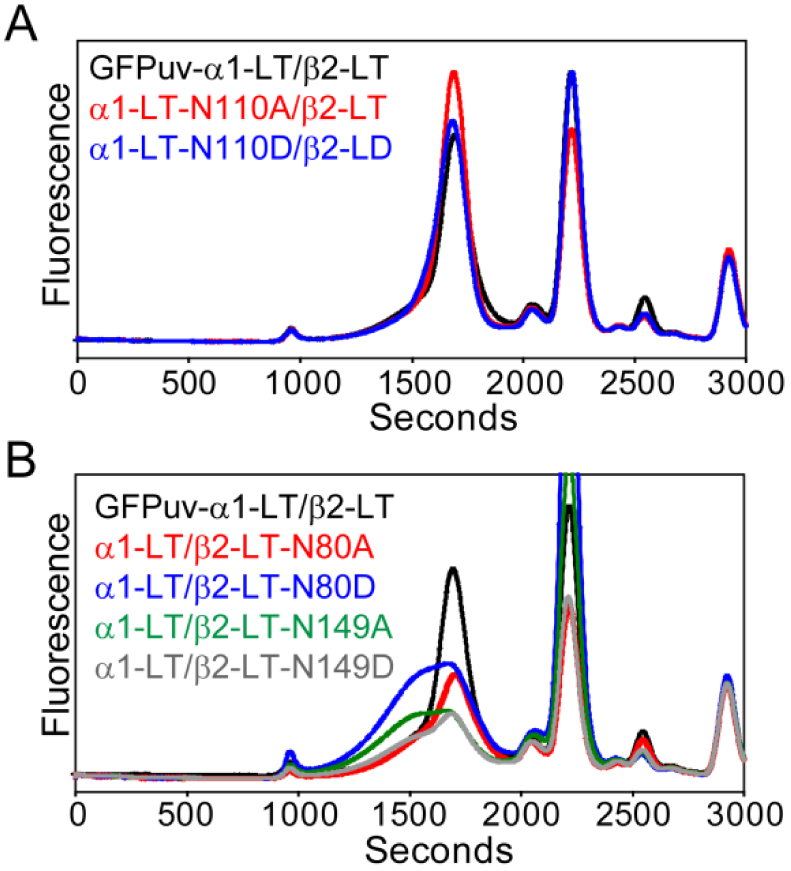
Mutation of other predicted glycosylation sites in the β2 subunit alters receptor expression and assembly. (A) Site directed mutation of a second consensus glycosylation site in the α1 subunit does not change receptor behavior, which is in contrast to mutation at two other glycosylation sites in the β2 subunit (B).

### Expression of the α/β GABA_A_ receptor in suspension culture

Encouraged by the positive screening results for the rat α1/β2 receptor in oocytes, we investigated suspension cell culture conditions to increase receptor yield for subsequent purification. Preliminary studies were performed in insect cells because homomeric GluCl and heteromeric GABA_A_ receptors have been expressed previously in Sf9 cells [23,64,65]. For these experiments, recombinant baculoviruses were created for the α1-EGFP and β2 (full length) subunits by shuttling the genes into separate pFastBac1 vectors, or combining the subunits into a single bicistronic pFastBac Dual vector (Fig 7A). Visible EGFP expression localized to the membrane was observed as early as one day post infection of Sf9 cells (Fig 7B). However, FSEC analysis indicated that the pentamer either failed to form properly into a homogeneous unit or was not maintained after whole cell solubilization with C_12_M at all tested time points (Fig 7C-D). Although we attempted to modify infection parameters, expression temperature, and detergent solubilization conditions, the α1/β2 pentameric receptor could not be isolated from Sf9 or High Five insect cells (S1 Fig).

**Fig 7.**
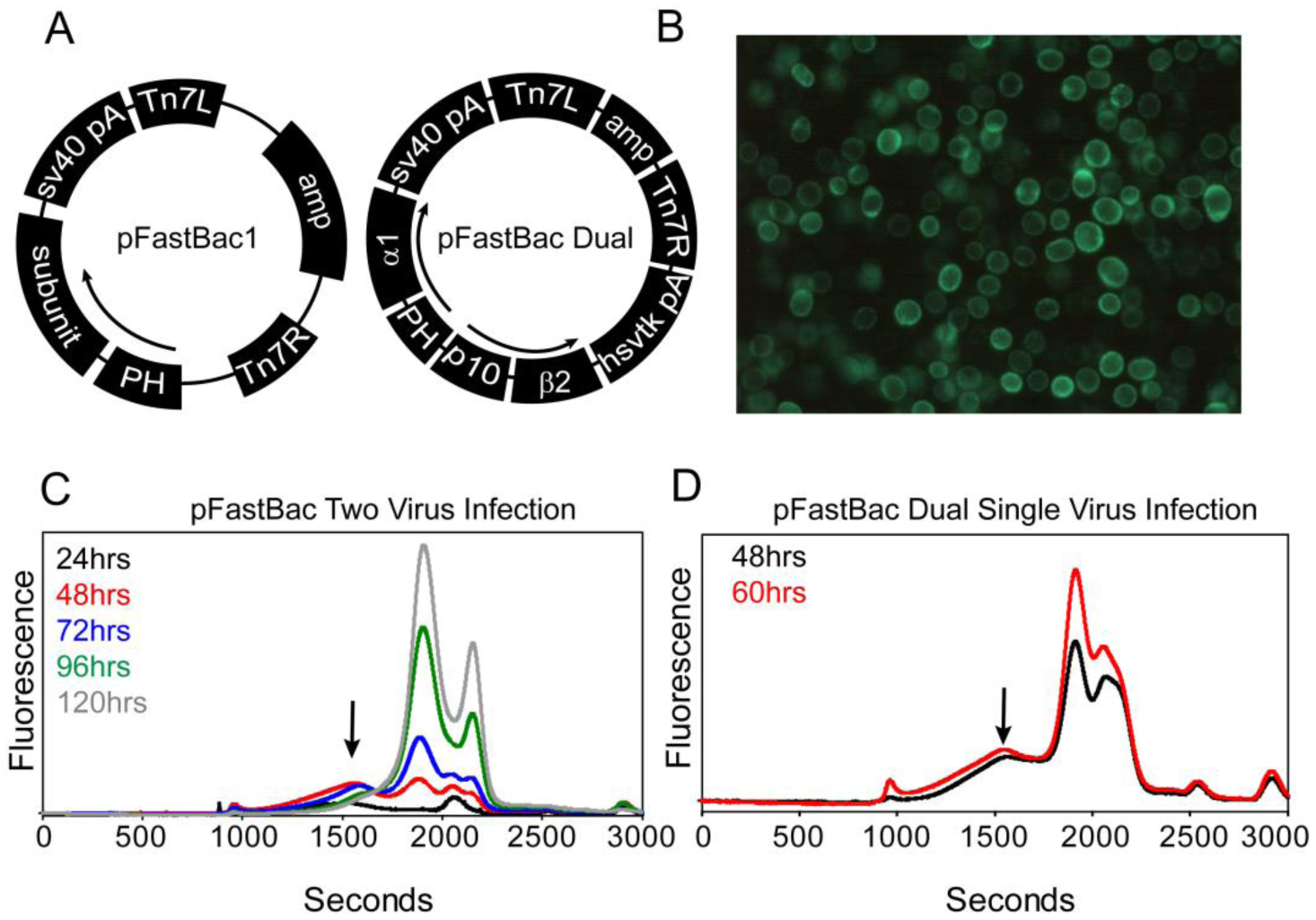
Pentameric α1-EGFP/β2 receptor cannot be isolated by detergent from Sf9 cells. (A) Diagram of insect cell expression vectors harboring one (pFastBac1) or both (pFastBac Dual) full length GABA_A_ subunits. (B) EGFP fluorescence is observed 48 hours post infection for pFastBac1 or pFastBac Dual constructs, suggesting receptor targeting to the cell membrane. (C) FSEC analysis at the indicated time points to measure expression suggests that most of the receptor breaks down upon whole cell solubilization with C_12_M detergent. (D) FSEC profiles do not improve using the bicistronic pFastBac Dual vector, which ensures each cell possesses a copy of both subunits.

We reasoned that mammalian cells may provide a more natural and stabilizing environment for heteromeric assemblies leading to successful membrane extraction. Therefore, the full length α1-EGFP and β2 subunits were cloned individually into a modified baculovirus vector, pEGBacMam [43], specifically designed to increase protein yields in adherent or suspension mammalian cells (Fig 8A). Efficacious virus was used to co-infect either TSA201 suspension cells or HEK293 suspension cells engineered to be deficient of *N*-acetylglucosaminyltransferase I (GnTI^−^) and thus eliminate complex *N*-glycan posttranslational modifications [66].

**Fig 8.**
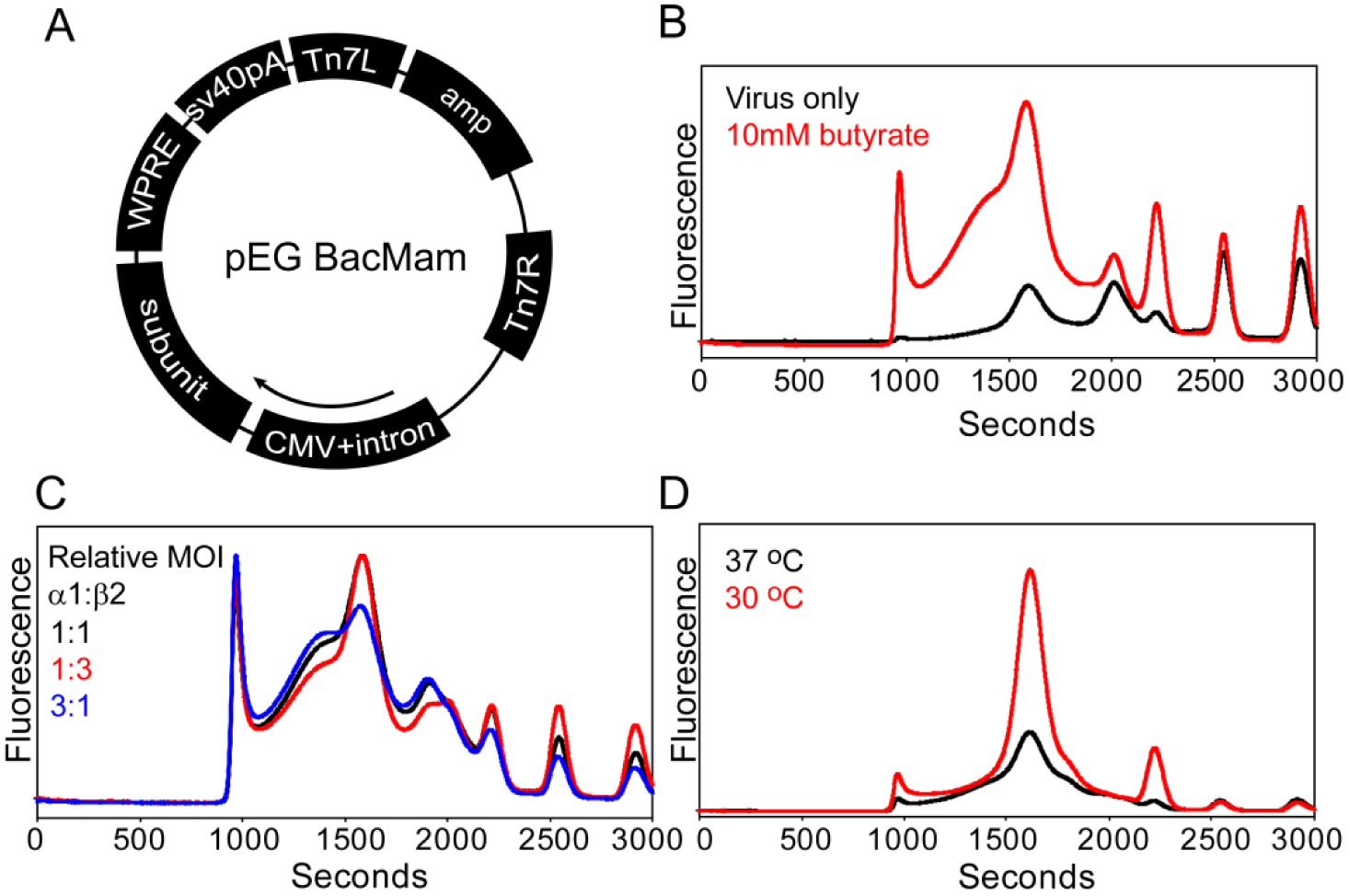
Identification of α1-EGFP/β2 receptor expression parameters in mammalian cells. (A) Diagram of pEGBacMam vector used for generating recombinant baculovirus to infect mammalian cells. (B) Addition of 10mM sodium butyrate to the virus transduced culture increases receptor yield but induces aggregation. (C) Infecting cells with a relative MOI≥1 (β:α) increases receptor monodispersity. (D) Dropping the temperature of infected cells increases receptor yield and reduces aggregation observed at higher temperatures.

In contrast to insect cells, the α1-EGFP/β2 pentamer was observed by FSEC analysis after whole cell solubilization with C_12_M 48 hours post infection (Fig 8B). Subsequently, expression of the α1-EGFP/β2 receptor was optimized by screening a variety of parameters for impact on fluorescence intensity and peak symmetry of the FSEC profile. Importantly, several of these parameters were found to be crucial to boosting receptor expression levels and behavior, as illustrated in Fig 8B-D. Addition of virus with a relative multiplicity-of-infection (MOI) ratio greater than one (β to α) increased receptor monodispersity, which may be a reflection of lower virulence for the β2 subunit. The inclusion of 10 mM sodium butyrate, a histone deacetylase inhibitor, 12 hours post infection increased expression levels ~5-fold relative to virus alone at the cost of increased propensity for receptor aggregation as seen by a strong leading shoulder in the elution peak. However, reducing culture temperature to 30 °C compensated for the negative impact of sodium butyrate at higher temperatures in addition to a concomitant three-fold increase in expression level observed at the 48 hour time point.

To assess the broad application of these expression conditions, other GABA_A_ receptor constructs were screened for changes in expression levels in mammalian cells. Indeed, these expression conditions worked well for substantially boosting expression of homomeric (ρ1-EGFP) and mutant heteromeric (α1-EGFP/β2-LT) ion channels (Fig 9A-B). Interestingly, exchanging the β2-LT subunit for the β1-LT isoform (similar M3/M4 loop truncation) further improved receptor homogeneity and enhanced expression by an additional three fold (Fig 9B). The increased expression levels allowed for an unambiguous evaluation of the importance of both β subunits to pentamer formation in mammalian cells. Consistent with observations made in oocytes, expression of α1-EGFP in the absence of either β subunit yields a polydisperse profile, indicating that assembly of a uniform pentameric receptor requires at least α and β subunits. Additionally, in the presence of either β subunit isoform, incorporation of the γ2S subunit to produce tri-heteromeric receptors reduced expression by more than 50% with these methods (S2 Fig). A time course of α1-EGFP/β1-LT receptor expression indicated that the receptor saturated 60-72 hours post infection (Fig 9C). Similar to α1-LT/β2-LT, the α1-LT/β1-LT receptor demonstrated agonist-driven channel activity in oocytes (Fig 9D).

**Fig 9.**
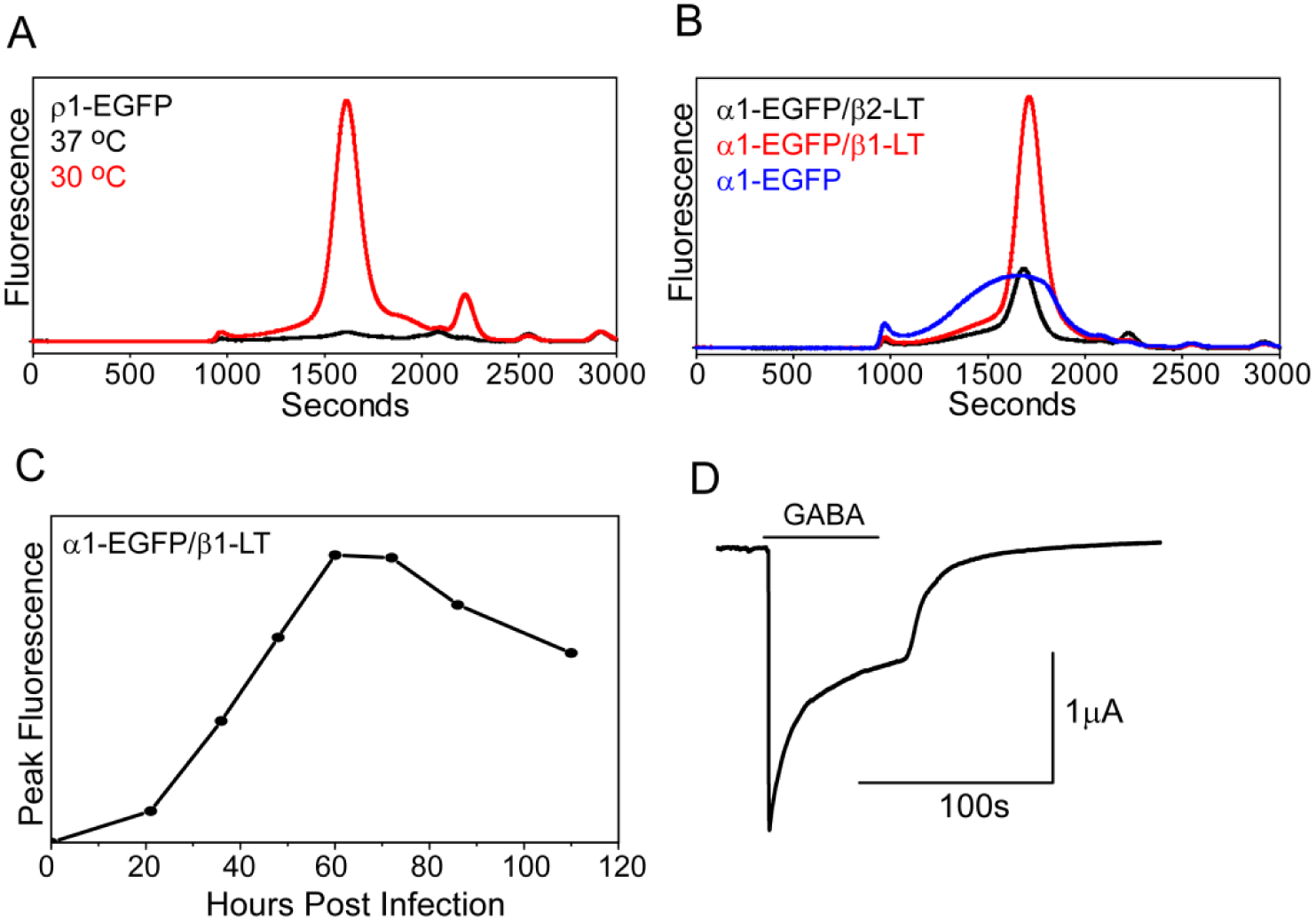
Optimized expression parameters apply to other receptor GABA_A_ receptor subtypes. (A) A boost in ρ1-EGFP expression levels is observed at low temperatures with addition of 10 mM sodium butyrate. (B) Similar to oocytes, the β subunit is required to form a homogeneous receptor in mammalian cells. In addition, the α1-EGFP/β1-LT receptor demonstrates higher expression levels than the α1-EGFP/β2-LT receptor. (C) Time course of α1-EGFP/β1-LT expression (relative MOI=0.6, α1:β1) followed by EGFP-FSEC suggests that the receptor demonstrates peak expression levels 60-72 hours post infection. (D) The α1-LT/β1-LT receptor displays GABA-gated current in oocytes by TEVC measurement.

### Purification and characterization of the α1/β1 receptor

Based on the expression studies outlined above, we pursued purification of the α1/β1 receptor combination from mammalian cell membranes. Preliminary experiments focused on the α1-EGFP/β1-LT receptor to facilitate optimization of purification strategies (S3 Fig). These experiments suggested that 50-55% of the expressed receptor was isolated from the membrane fraction based on EGFP-FSEC analysis. Subsequently, other receptor constructs containing α1 subunits devoid of EGFP or with truncated M3/M4 loop were isolated and characterized through ligand binding assays. In general, a simple two-step protocol was chosen that combines immobilized metal affinity chromatography (IMAC) followed by gel filtration to remove aggregates and degradation products. An octa-histidine tag was fused to the C-terminus following the thrombin protease recognition sequence in either one or both subunits to facilitate IMAC purification and ligand binding analysis via scintillation proximity assay (SPA). Other elements of primary sequence, such as glycosylation sites, were retained in both subunits to maximize yields.

Following solubilization with 40 mM C_12_M, the α1/β1 receptor was purified successfully into either 1 mM C_12_M or 0.3 mM lauryl maltose neopentyl glycol (L-MNG) detergent micelles (Fig 10). Typical yields from IMAC purification for all constructs were in the range of 0.4-0.7 mg/L (~2.5 nmol/L) of suspension culture based on absorbance at 280 nm assuming an average 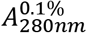 (Materials and Methods). Isolation of monodisperse receptor was obtained from preparative size exclusion chromatography (Fig 10A-B). SDS-PAGE analysis indicated that the mobility of each subunit corresponds to the expected molecular weight (α1-LT=43 kDa; β1-LT=42 kDa; Fig 10C). Confirmation of the band identities was determined by LC-MS. In addition, each subunit ran as diffuse band(s) that would collapse upon treatment with an endoglycosidase cocktail (S4 Fig), suggesting that the receptor is glycosylated.

**Fig 10.**
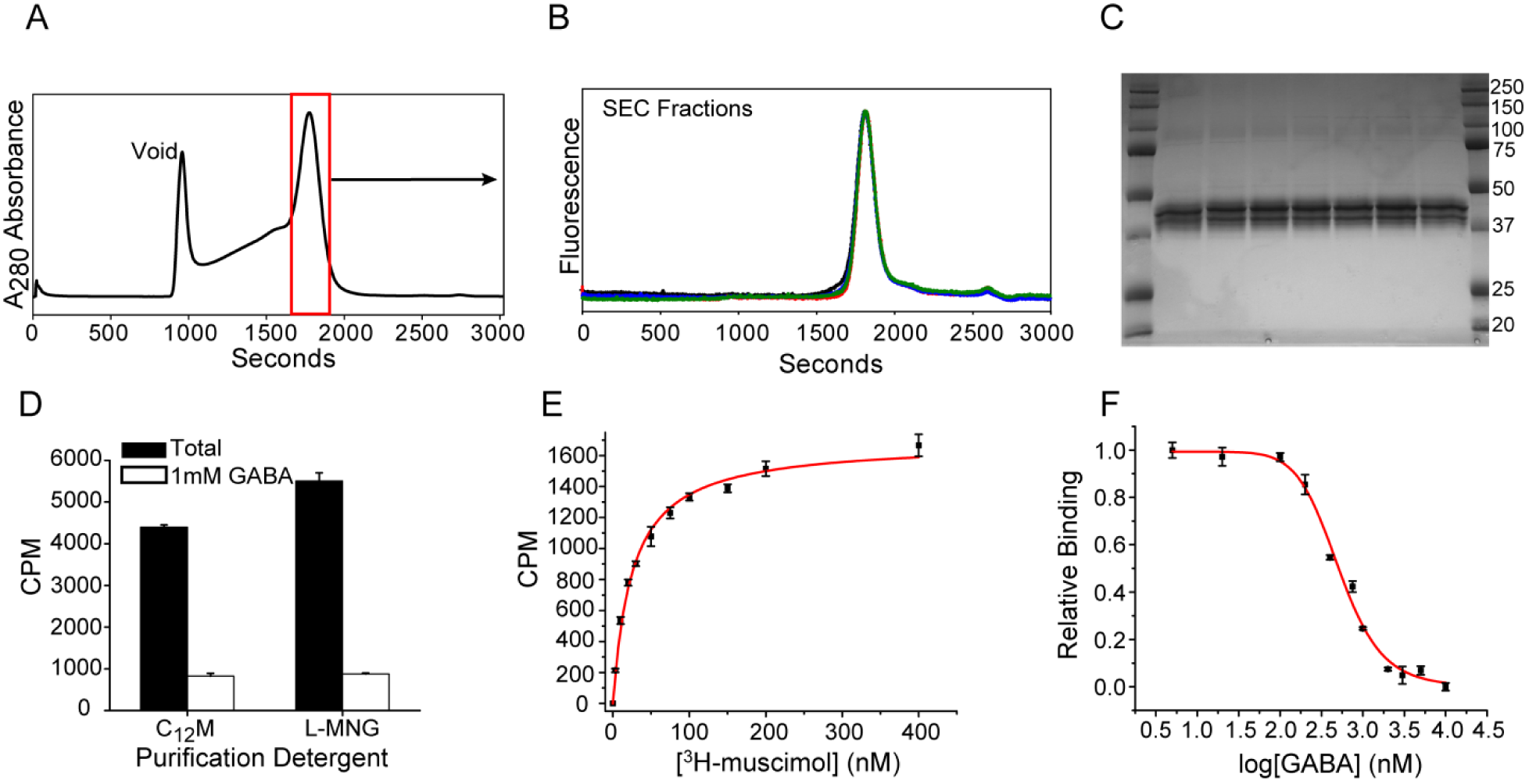
Purification and characterization of the α1/β1 receptor. (A) Preparative size exclusion chromatography of IMAC-purified material isolates homogeneous receptor as observed by FSEC analysis of the elution fractions (B). (C) SDS-PAGE analysis indicates that purified receptor shows diffuse subunit bands, consistent with glycosylation. (D) Purified receptor shows specific binding in C_12_M or L-MNG detergent. Ligand binding experiments by SPA reveal high affinity binding sites for muscimol (E) or GABA (F) as measured by a direct binding isotherm or competition assay, respectively. Results of the fits are highlighted in the main text.

The integrity of purified receptor was investigated by measuring ^3^H-muscimol binding activity using SPA. Specific binding of ^3^H-muscimol was observed in both C_12_M and L-MNG (Fig 10D). Titration of ^3^H-muscimol produced a monotonic binding isotherm (Fig 10E). Non-linear least squares fitting assuming a single site binding model revealed a *K*_D_ of 24±1.1 nM, consistent with previously reported values for the α1/β1 receptor in intact cells [67]. Furthermore, a competition assay in which 100 nM ^3^H-muscimol was displaced by increasing concentrations of GABA demonstrated an IC_50_ of 507±28 nM and a *K*_i_ of 98±6.5 nM (Fig 10F). These results are consistent with the presence of high affinity agonist binding sites formed by the interface of α1 and β1 subunits.

## Discussion

Here we have shown that construct screening in *Xenopus* oocytes in combination with FSEC is a viable alternative to traditional approaches, such as *in vitro* transient transfection, to identify a suitable heteromeric GABA_A_ receptor for large-scale expression and purification. This method presents a number of advantages over plasmid DNA transfection, such as straightforward gene delivery, dosage control and ease of manipulation. To enhance plasmid transfection efficiencies, cellular endocytosis is facilitated by cationic “carrier” molecules such as charged lipids or small chemical compounds like calcium phosphate. However, transfection reagents can induce a cytotoxic response and the relative ease of transfection varies with the cell line which can produce artificially low gene expression levels. Furthermore, protocol optimization becomes more imperative for multi-gene studies, such as with GABA_A_ receptors, in which overall transfection efficiency becomes compounded. Although viral transduction is an efficient strategy for delivering genetic cargo, making a library of high titer recombinant viruses for expression studies can be time consuming, costly and counterproductive for the iterative pathway of identifying a lead gene construct for expression and downstream structural studies.

Screening GABA_A_ receptor genes in oocytes not only led to identification of the α/β subunit combination as an excellent candidate for further investigation, but also informed gene design since expression levels and homogeneity were modulated by construct mutagenesis. The application of the FSEC technique to facilitate oocyte screening adds a new dimension to the process of identifying the best expressing and behaving constructs. Oocytes provide a tractable platform for identifying functional constructs, but investigating electrophysiological properties alone may lead to false positives since large expression levels are not needed to generate ion channel currents in oocytes. For instance, α1/β2 and α1/β2/γ2S will form functional channels, but the tri-heteromeric receptor demonstrated substantially reduced expression levels. Importantly, the oocyte expression profiles of receptor subtypes investigated here were mirrored by mammalian cells, suggesting continuity between disparate expression systems.

Combining construct screening in oocytes with FSEC analysis has uncovered important determinants for receptor assembly. Studies have suggested that high affinity binding sites in GABA_A_ receptors are formed only in the presence of the β subunit [68-70]. As shown in oocytes and mammalian cells, the requirement of the β subunit to form a functional channel is visualized at the level of subunit association. In the absence of the β subunit, a homogeneous α1 pentamer fails to form leading to broad, polydisperse gel filtration profiles that are the hallmark of destabilized oligomers. Furthermore, the observation that a monodisperse pentamer requires the β subunit was validated by FRET and electrophysiology experiments in oocytes. The lack of an FSEC-defined receptor composed of α1 only or α1/γ2S correlates with the lack of functional surface expression previously seen in oocytes [71] or in transfected L929 cells [72]. These experiments also demonstrate that large truncations in the cytoplasmic M3/M4 loop are not detrimental to expression, subunit association or macroscopic gating properties, which is consistent with previous studies of homomeric serotonin and ρ1 receptors [57]. In contrast, interference with subunit glycosylation had a substantial impact on expression and assembly.

To our surprise, the most tractable subunit combination identified in oocytes, α1/β2, could not be isolated with detergent from insect cells. A number of studies have shown that di- and tri-heteromeric GABA_A_ receptors, including α1/β2, can be expressed as functional channels with high affinity ligand binding sites in Sf9 cells using recombinant baculovirus [64,65,69]. Notably, these studies examined receptor activity in cell membranes or intact cells. We found that the α1-EGFP/β2 combination expressed to the membrane, which suggested that the receptor is being properly secreted and targeted. However, detergent solubilization resulted in gross breakdown of the receptor as judged by FSEC analysis, which indicated that insect cells are not a practical means of over-expressing and purifying large quantities of this receptor. Given the success of our laboratory with GluCl in Sf9 cells, this observation may illustrate a distinguishing feature between stable assembly or maintenance requirements of homo- and heteromeric Cys-loop receptors.

Although the α/β receptor could be expressed in mammalian cells, high level expression was only obtained after exploring culture conditions and baculovirus infection parameters. Remarkably, the addition of 10 mM sodium butyrate and the reduction of culture temperature to 30°C, both of which have been shown to strongly impact protein expression in mammalian cells [47,73,74], demonstrated a synergistic effect on receptor expression and behavior. For instance, the sodium butyrate-induced protein aggregation observed during expression at 37 °C was substantially reduced when the culture temperature was shifted to 30 °C.

In addition to culture conditions, co-infection with the β1 subunit drove expression levels even higher than with the β2 subunit. The origin of the difference in expression levels between α1/β1 and α1/β2 subtypes is not clear. Even though the β1 subunit shares 77% sequence identity with the β2 subunit, it is the most divergent of the three known β isoforms. Because most of the sequence variation in the mature subunit arises from the M3/M4 loop which can be removed without impairing expression or function based on these studies, increased expression levels of the α1/β1 receptor relative to α1/β2 may be confined to ~10% of sequence that is not strictly conserved. A systematic examination of signal peptide strength and mutagenesis of non-conserved residues would be required to elucidate the impact of β subunit identity on receptor expression. Despite the determined optimal conditions, expression of α/β/γ2S was still attenuated relative to α/β, which may be a consequence of infection with multiple viruses that is detrimental to cell health. The pliability of pEG BacMam to combine multiple genes into a single vector could be a possible means of increasing α/β/γ2S expression levels by eliminating the need for infecting with multiple viruses.

The yields of α1/β1 receptor obtained from expression and purification of mammalian cell membranes are comparable with that reported for human α1/β3 with an amino-terminal FLAG tag from an HEK293-derived tetracycline-inducable stable cell line [75]. The benefit of the baculovirus system as described here is the ability to rapidly and efficiently express many different constructs without the time consuming process of generating stable cell lines, which is highly advantageous for studies that seek to unveil the role of specific residues in protein structure and function through site directed mutagenesis. Most importantly, the receptor extracted from the membrane maintains high affinity ligand binding capability, suggesting a defined subunit organization that represents native associations.

In conclusion, the methods described here have garnered exceptional results toward efficiently expressing and purifying a functional heteromeric GABA_A_ receptor for structural analysis. Although this approach is focused uniquely upon GABA_A_ receptors, we anticipate that these methods will be broadly applicable to other complex protein systems that may require multiple protein subunits or binding partners. In general, expression of other eukaryotic membrane proteins in our laboratory has benefited immensely using the mammalian system described here [76-81].

## Acknowledgements

The authors wish to thank Dr. David S Weiss of Baylor College of Medicine for providing the α1, β2 and γ2S genes in the pGEM vector. We thank Dr. David C Dawson for providing oocytes and Dr. Haining Zhong for the mKalama fluorescent protein. We also thank Dr. Stefanie Petrie of the OHSU Advanced Light Microscopy Core for expertise and data collection with the Zeiss LSM710 laser scanning confocal microscope.

## Supporting Information

**S1 Fig.**
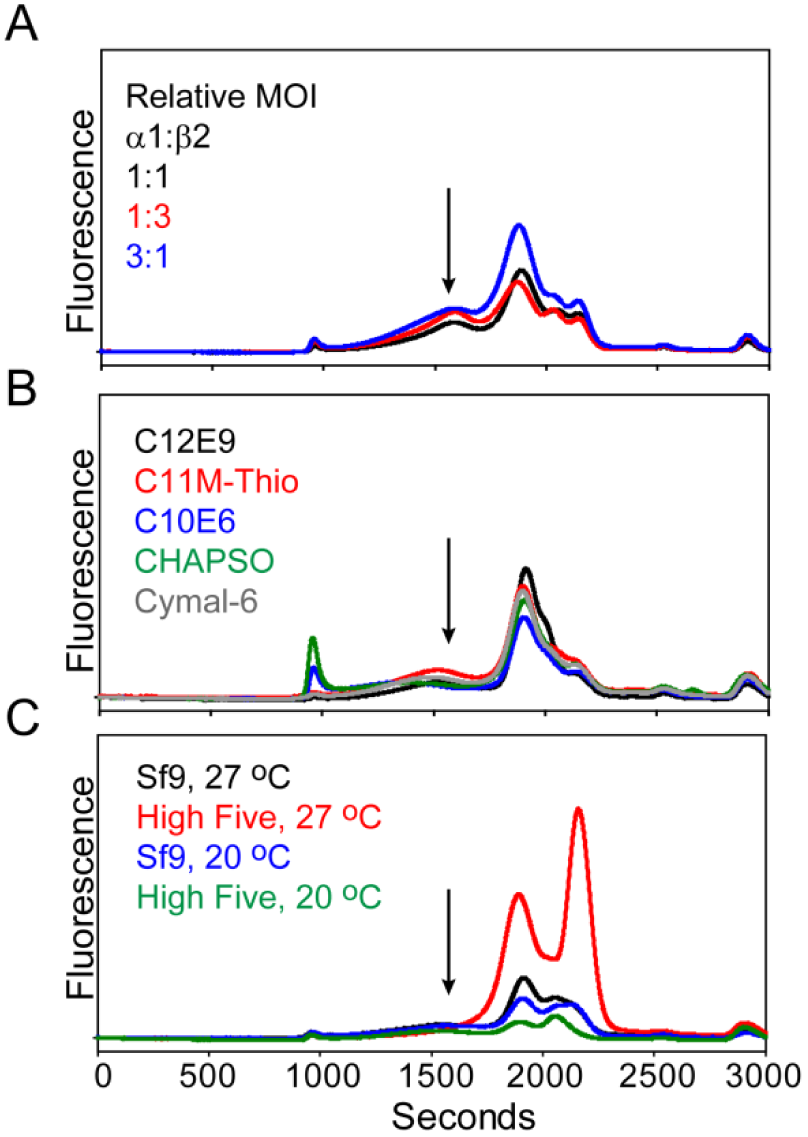
Screening for conditions to express and extract α1-EGFP/β2 from insect cells. (A) Infection of Sf9 cells with two separate baculoviruses with different ratios (relative MOI) does not improve expression as observed 72 hours post infection. (B) Whole-cell solubilization with a panel of detergents does not prevent receptor breakdown. (C) Infection of High Five cells or shifting cultures to lower temperatures does not improve receptor recovery. The traces are plotted on the same scale as Fig 7C.

**S2 Fig.**
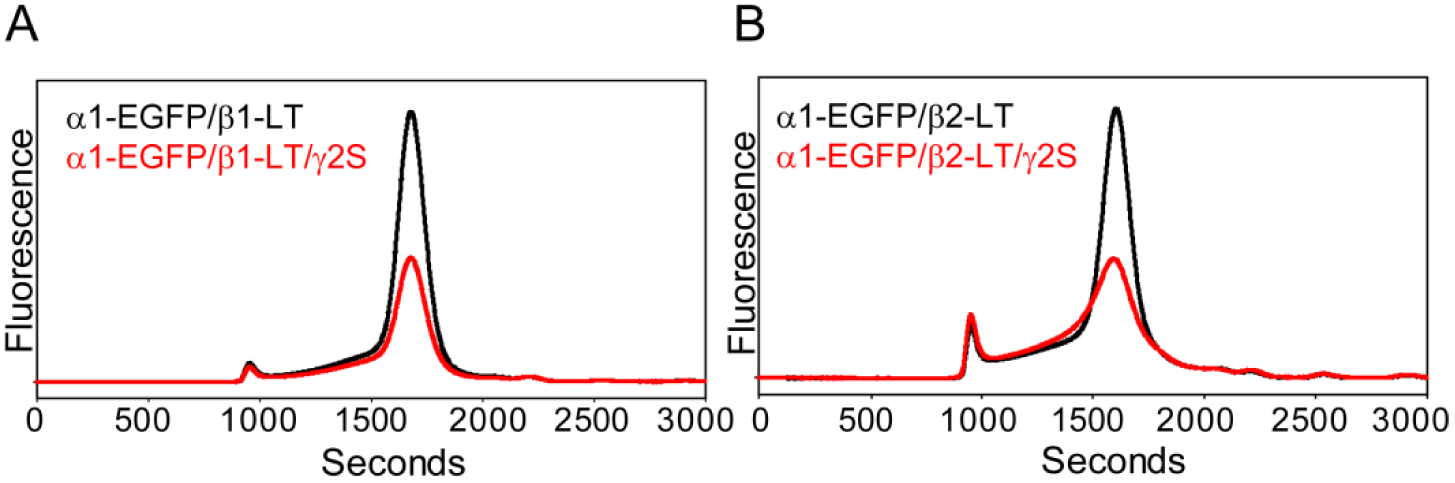
Expression of tri-heteromeric GABA_A_ receptors in mammalian cells. Incorporation of the γ2S subunit substantially reduces receptor expression levels relative to α1-EGFP/β1-LT (A) or α1-EGFP/β2-LT (B) at similar time points.

**S3 Fig.**
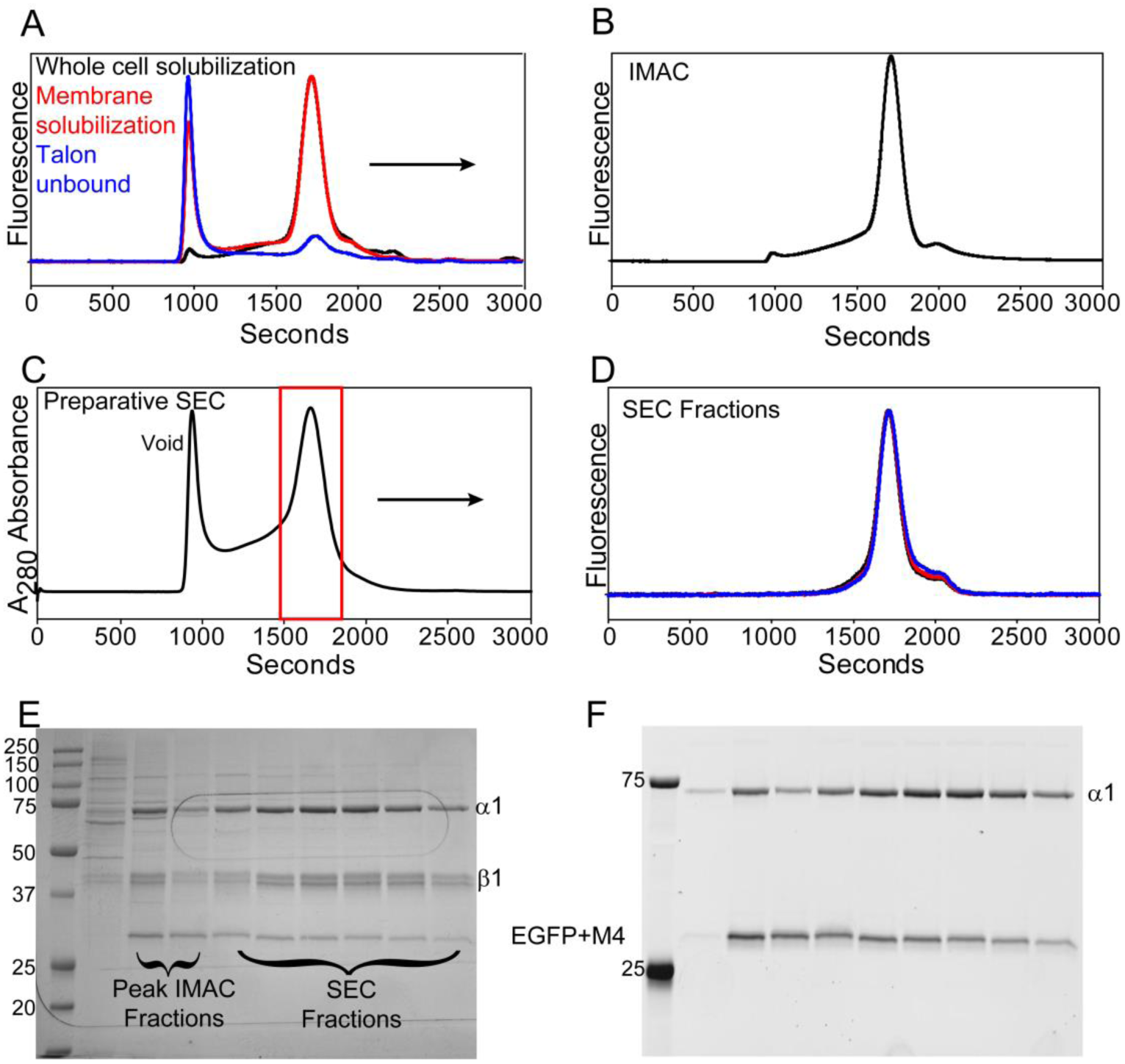
Purification of α1-EGFP/β1-LT. (A) The receptor demonstrates a similar FSEC profile whether solubilized from whole cells (black trace) or from the membrane fraction (red trace). The traces are normalized to emphasize similarities. Based on the FSEC analysis, approximately 50-55% (~3.3 mg) of the expressed receptor (6.4 L culture) was extracted from the membrane fraction. Of this material, 90% of the receptor was bound to Talon resin after three hours batch binding as reported by a depletion of receptor from the solution (blue trace). Approximately 3mg of receptor (0.5 mg/L or ~2 nmol/L) eluted from the resin (B), and was concentrated for preparative size exclusion chromatography (C). Peak fractions as identified by the red box in (C) were analyzed by FSEC for homogeneity (D). SDS-PAGE analysis demonstrates the purity of the receptor as a function of purification steps. The first lane shows contaminants eliminated from a low concentration of imidazole (50 mM) in the buffer wash. Notably, the presence of EGFP in the α1 M3/M4 loop enhanced proteolytic cleavage relative to non-fusion constructs, giving rise to a ~30 kDa band in SDS-PAGE analysis (E). Confirmation that this band contained EGFP was made by in gel fluorescence prior to fixing and staining (F). Although we did not explicitly determine the identity of this band, the migration position is consistent with EGFP plus the M4 α-helix. However, cleavage of the loop likely does not disrupt native associations since the purified receptor demonstrated monodispersity by FSEC analysis as shown in panel D.

**S4 Fig.**
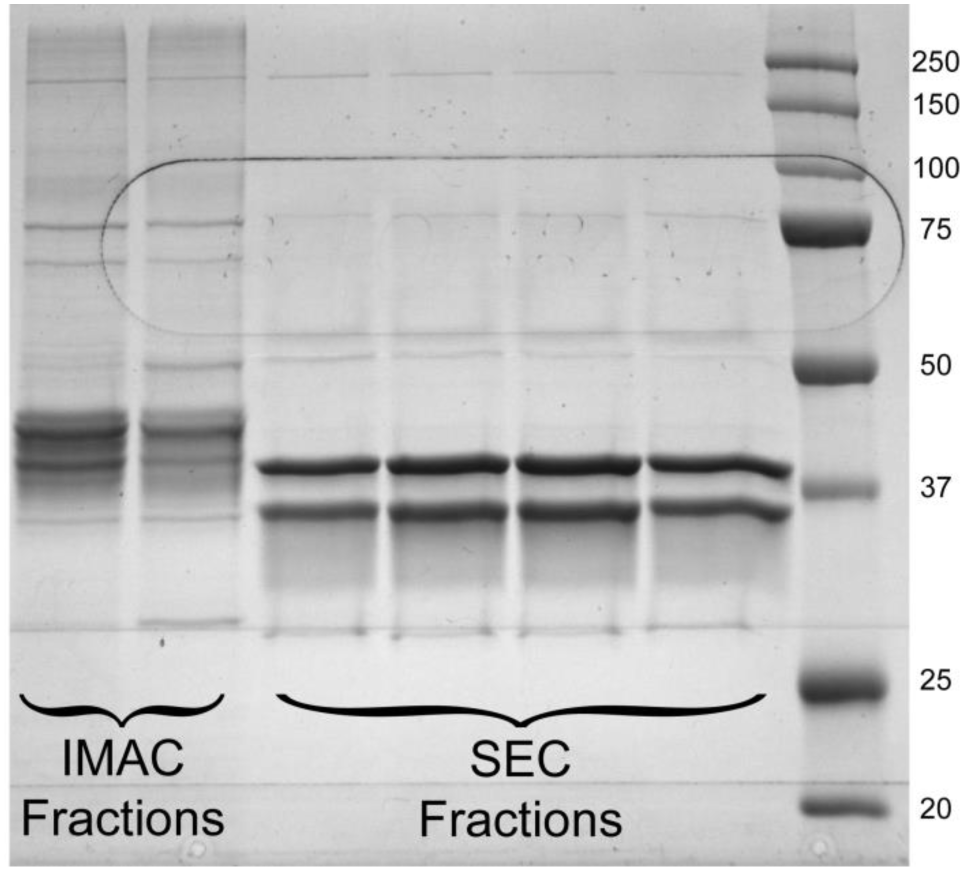
Endoglycosidase treatment removes N-glycans from α1 and β1 subunits. Following IMAC purification, the receptor was treated with EndoH and EndoF_3_ for two hours at room temperature. After purification by size exclusion chromatography, the receptor was analyzed by SDS-PAGE and showed that treatment caused the diffuse bands seen in the IMAC fractions to collapse into a single prominent band for both subunits.

